# From video to encounter histories: individual identification and machine vision for salmonid capture–recapture monitoring

**DOI:** 10.64898/2026.09.03.749040

**Authors:** Konrad Karlsson

## Abstract

Individual encounter histories are central to capture–recapture models, but fisheries monitoring is often reduced to counts that do not account for variation in detection probability. This study presents a machine-vision workflow for converting long-term video surveillance of Atlantic salmon (*Salmo salar*) and sea trout (*Salmo trutta*) spawning runs into individual-level data for capture–recapture analysis. The workflow detects fish-positive frames, saves video clips, extracts fish-head regions of interest, and organizes cropped images for re-identification. A binary EfficientNetB0 fish detector achieved validation accuracy of 0.9918 and PR AUC of 0.9992; at a conservative threshold of 0.98, validation false positives were eliminated while retaining 94.3% recall. A YOLOv8n model localized fish-head regions, and an EfficientNetB0 ArcFace model trained on head images from 700 identities achieved 99.80% accuracy among accepted known matches and an image-weighted false-accept rate of 0.81% when non-training identities were treated as unknown. Zero-shot closed-set retrieval achieved 99.17% top-1 accuracy across 1,416 identities excluded from model training, demonstrating strong generalization to previously unseen identities.

## 1 Introduction

Individual identification provides fundamental data for many hierarchical models used in fish and wildlife management (Kéry and Royle, 2016, 2021). Many populations of Atlantic salmon (*Salmo salar*, hereafter salmon) and sea trout (*Salmo trutta*) are vulnerable (HELCOM, 2011), and estimates of spawning-run size therefore need to be as accurate and precise as possible. In fisheries monitoring, however, individuals are often not explicitly identified, and detections or captures are instead summarized as counts. If counts are higher in one year than another, or at one location than another, this may indicate an increasing population or higher local abundance, all else being equal (e.g. Roser et al., 2025). However, the assumption that all else is equal is rarely realistic. Changes or differences in detection probability can strongly influence relative-abundance estimates and may even mask population declines (Erisman et al., 2011; Charbonneau et al., 2025; Karlsson et al., 2025). Relative abundance is often expressed as the number of detected or captured fish, *n*, whereas population size, *N*, depends on both the number detected and the probability of detection, *p*. In its simplest form, the expected count is *n = Np*, and population size can therefore be estimated as *N = n/p* when detection probability is known or estimated (Kéry and Royle, 2021). Population-size estimation therefore explicitly incorporates the observation process, allowing counts to be interpreted relative to detection probability rather than assuming that observed counts directly represent true abundance.

Capture–recapture models explicitly estimate detection probability, *p*, from individual encounter histories, that is, records of individual detections and non-detections across sampling occasions (Otis et al., 1978). In fish studies, individuals are traditionally identified using artificial markers, including internal tags such as passive integrated transponder (PIT) tags (Raabe et al., 2014), acoustic transmitters (Kazyak et al., 2020), or external T-bar tags (Karlsson et al., 2025). In contrast, many large mammals are identified non-invasively from photographs, using unique natural markings as individual identifiers (Würsig and Jefferson, 1990; Karanth, 1995; Royle et al., 2018). One reason for this difference may be that fish populations, even when threatened, often contain many more individuals than populations of large mammals, while identification from natural markings has traditionally required manual comparison of images. However, photo-based identification has been applied to smaller fish populations, for example in a pike (*Esox lucius*) study where 39 individuals were identified from 66 captures using natural markings (Karlsson and Kari, 2020). Manual photo-identification does not scale easily to larger datasets. For example, the International Whaling Commission noted that matching a single whale photograph against a catalogue of 800 individuals could require approximately 8 hours, which was considered an upper practical limit for manual identification (Würsig and Jefferson, 1990).

Recent advances in deep learning–based computer vision, supported by large labelled image datasets, convolutional neural-network architectures, and accessible software tools, have made automated processing of ecological image and video data increasingly practical (Deng et al., 2009; Krizhevsky et al., 2012; Chollet, 2017; Weinstein, 2018; Christin et al., 2019). These developments now support animal detection, classification, and individual re-identification in large image datasets (Steenweg et al., 2017; Norouzzadeh et al., 2018; Schneider et al., 2019; Vidal et al., 2021; Yesharim et al., 2026). At the same time, camera-based monitoring is increasingly used in fish ecology, including both manually reviewed underwater video surveys and automated machine-vision approaches for detecting and monitoring fish (Soom et al., 2022; Atlas et al., 2023; Karlsson, 2024; Bowman et al., 2026).

However, most fish-monitoring applications still summarize observations as detections, counts, or relative-abundance indices (Erisman et al., 2011), rather than retaining individual-level information that can be used directly in capture–recapture models. A natural next step is therefore to link machine-vision detection with individual identification, so that video data can be transformed into encounter histories for capture–recapture analysis. This study presents a machine-vision pipeline consisting of several linked models that: (i) detect fish-positive frames and save video clips of fish passages, (ii) extract regions of interest (ROIs) containing the visual information used for individual identification, and (iii) organize ROIs from each video clip into folders for downstream individual-identification analysis.

This machine-vision pipeline was developed for monitoring salmon and sea trout spawning runs, which may extend from early summer through autumn (Klemetsen et al., 2003; Weedop et al., 2026). The monitoring design was motivated by capture–recapture approaches previously used to estimate salmonid abundance during migration (Schwarz and Dempson, 1994; Mäntyniemi and Romakkaniemi, 2002) and by the more general multinomial framework for modelling abundance from individual encounter histories described by Kéry and Royle (2016). In a two-station design, fish detected and identified at a downstream camera station can subsequently be re-identified at an upstream station, producing encounter histories analogous to those generated in conventional capture–recapture studies. These repeated detections can be used to estimate detection probability and abundance without physically capturing or marking fish. Although related two-station capture– recapture designs have primarily been applied to juvenile salmonids (smolt) migrating downstream to sea, the same logic can be applied to adults migrating upstream to spawning grounds.

## 2 Materials and methods

### 2.1 Experiments involving animals

The study used video material of fish collected by non-invasive camera monitoring in their natural environment. No fish were captured, handled, marked, transported, experimentally manipulated, or euthanized for this study. Therefore, animal ethics approval was not obtained because the work was based solely on observational video data and involved no physical intervention with animals. Fieldwork permits were not required because fish were recorded without manipulation of their environment or movements. Camera deployments at all sites were discussed with the relevant fisheries managers.

### 2.2 Video data sources and recording configurations

Video recordings were obtained using two camera configurations. In the first configuration, underwater cameras were placed directly on the riverbed. This setup was used in Dalälven, Hartijoki, Mörrumsån, and Skeboån. In the second configuration, an IP camera mounted in a room recorded fish through a glass wall in a fish passage in Stornorrfors, Vindelälven. These recordings provided the source material for training and evaluating the fish-presence detector, fish-head region-of-interest detector, and individual re-identification model. The underwater recordings were collected using IP underwater cameras with long PoE cables connected to a network video recorder (NVR; Karlsson, 2024). The NVR stored continuous video and retained auxiliary metadata, including camera identity and date-time information. Video quality was set to retain sufficient detail for individual identification, with recordings generally made at high-definition resolution. Infrared illumination was used during low-light conditions when available. The IP cameras in Stornorrfors recorded at near-4K resolution because fish were farther from the camera and the full body needed to be visible in one frame. The passage tunnel was continuously illuminated with artificial light.

### 2.3 Machine vision pipeline

#### 2.3.1 Fish-presence detection model

The first stage of the image-processing pipeline consisted of a binary classifier that determined whether a frame contained a fish. This detector was used as a screening model before subsequent region-of-interest cropping and individual-identification stages. The model produced a single sigmoid output, consistent with standard probabilistic binary classification (Bishop, 2006). The class “no_fish” was encoded as 0 and “fish” as 1.

Image data were constructed from network video recordings from five locations and four years, in total eight combinations (Figure 1). The fish-detector dataset contained 68,489 images, comprising 24,685 “fish” and 43,804 “no_fish” images. Of these, 54,790 images were used for training, comprising 19,747 “fish” and 35,043 “no_fish” images, and 13,699 images were used for validation, comprising 4,938 “fish” and 8,761 “no_fish” images. Prior to model training, the dataset was divided into training and validation subsets using an 80:20 split within each combination of class and location-year group.

**Figure 1.**
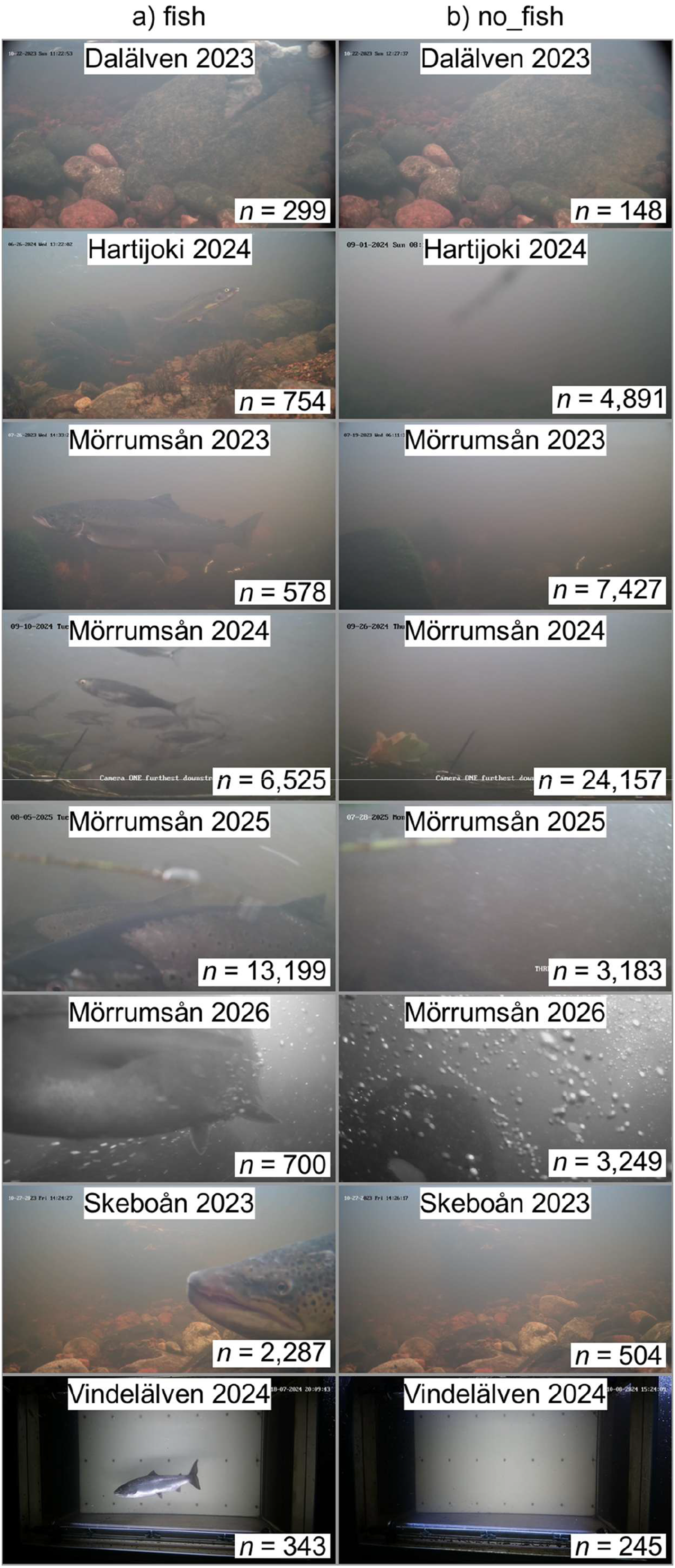
Sample images for the fish detector model, with one image from each class-location-year combination. Images labelled “fish” are shown in column a), and images labelled “no_fish” are shown in column b). The number of images in each combination is printed in the corresponding image.

The model was implemented in TensorFlow/Keras (Abadi et al., 2016; Chollet, 2015) using an EfficientNetB0 (Tan and Le, 2019) backbone initialized with ImageNet weights (Deng et al., 2009). The EfficientNetB0 backbone was used without its original classification head and was kept frozen during training. Only the newly added binary classification head was trained. Input images were resized to 400 × 224 RGB pixels. During training, mild image augmentation was applied, including random horizontal flipping, small random rotations, and random contrast variation.

The classification head consisted of global average pooling, a fully connected dense layer with 256 units and ReLU activation, dropout with a rate of 0.5 (Srivastava et al., 2014), and a final sigmoid output node named fish_probability. The final output layer was kept in float32 precision to maintain numerical stability when mixed-precision training was enabled (Micikevicius et al., 2018). Training was performed using binary cross-entropy loss and the Adam optimizer (Kingma and Ba, 2015) with an initial learning rate of 1 × 10^−4^. The model was trained for up to 15 epochs with a batch size of 8. Mixed-precision training was enabled when a compatible GPU was available. Class weights were calculated from the number of training images in the “fish” and “no_fish” classes to compensate for class imbalance. The training dataset was shuffled, whereas the validation dataset was evaluated without shuffling.

Model selection was based on validation precision–recall area under the curve (PR AUC), which is useful for evaluating binary classifiers when class balance and false-positive control are important (Saito and Rehmsmeier, 2015). During training, the best-performing model was saved using model checkpointing. Early stopping was applied with restoration of the best weights, and the learning rate was reduced when validation PR AUC stopped improving. Training history, model checkpoints, training curves, and evaluation summaries were saved automatically.

After training, the model was evaluated on the validation set. Standard classification metrics were calculated, including accuracy, precision, recall, specificity, F1 score, balanced accuracy, false-positive rate, false-negative rate, PR AUC, and receiver operating characteristic (ROC) AUC (Fawcett, 2006). Threshold-dependent performance was then evaluated by applying fish-probability decision thresholds from 0 to 1 in increments of 0.001. For each threshold, the numbers of true positives, true negatives, false positives, and false negatives were recorded, and precision, recall, specificity, and F1 score were calculated. Deployment-relevant high-confidence thresholds of 0.90, 0.95, 0.97, 0.98, and 0.99 were summarized separately to assess the trade-off between reducing false-positive fish detections and retaining true fish detections.

#### 2.3.2 Fish-head ROI detection model

The fish-head detection dataset contained 25,613 images, split into 20,490 training images and 5,123 validation images. Across the dataset, 7,180 fish-head bounding boxes were manually annotated. Images without annotated head boxes were retained as negative/background images when no head was visible in sufficient detail for individual identification (Figure 2). A single-class object-detection model was trained to localize fish heads for subsequent region-of-interest extraction and individual-identification analysis. The detector was implemented using YOLOv8n (Redmon et al., 2016; Jocher et al., 2023) and initialized from pretrained weights. All annotated head boxes were treated as a single class. Images were resized to 640 pixels during both training and validation. Training was performed on GPU using mixed precision (Micikevicius et al., 2018), a batch size of 16, and RAM image caching. The model was trained for a maximum of 70 epochs, with early stopping after 20 epochs without validation improvement. The full network was trainable, with no frozen layers, and reproducibility was controlled using a fixed random seed.

**Figure 2.**
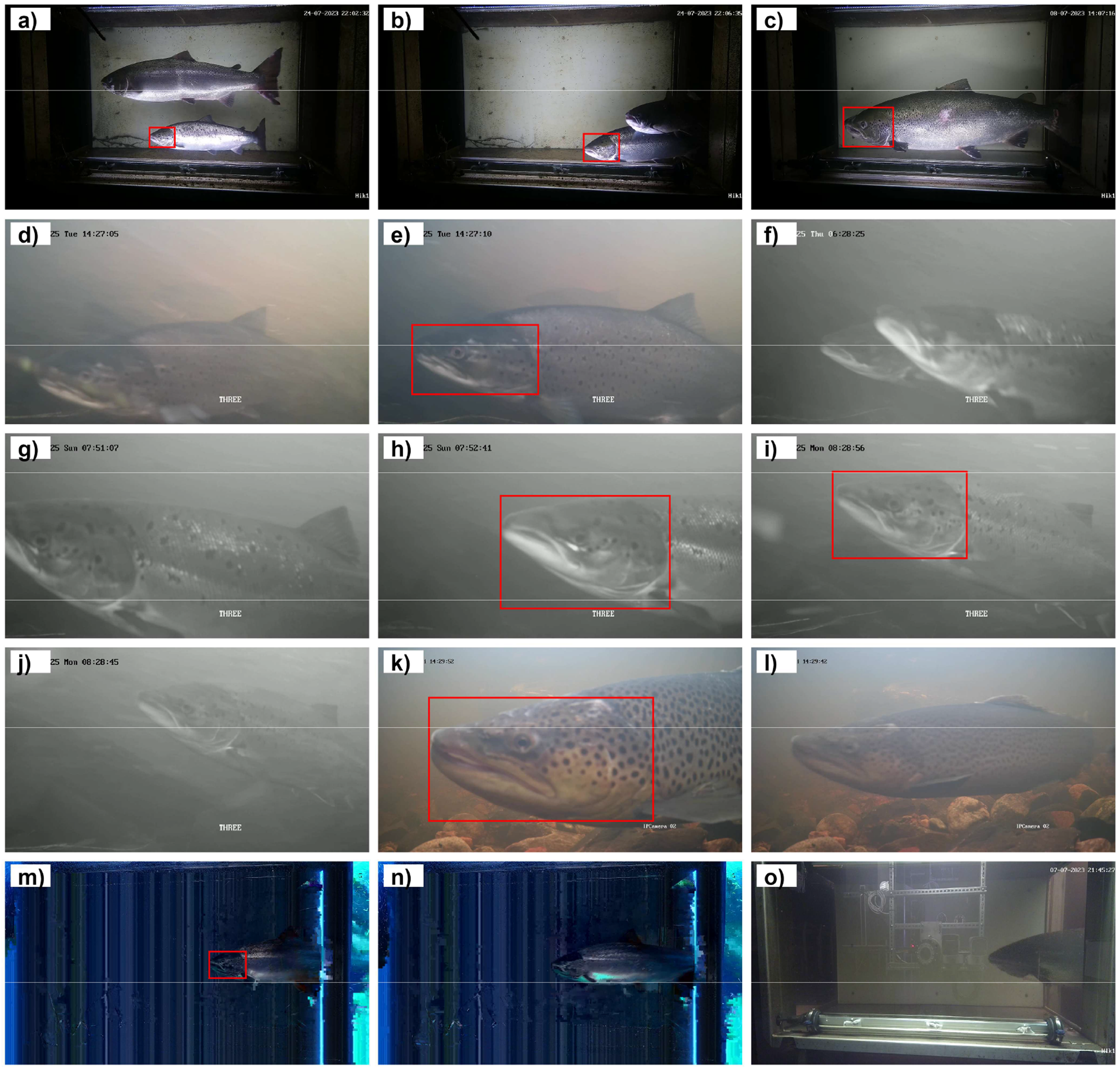
Examples of image annotations used for training and validation of the YOLOv8n fish-head detector. The figure shows 15 selected images illustrating the annotation procedure used to define fish-head regions of interest for subsequent cropping and individual identification, panels a) to o). The images and red bounding boxes are part of the manually labelled training and validation data and therefore represent human-defined annotations, not model predictions. Fish heads were annotated when they were visible in sufficient detail to be considered useful for individual identification; otherwise, no annotation was added. Partial or insufficiently visible heads, such as the example in g), were not annotated. Images a) to c), and m) to o) are from Vindelälven and were filmed through a glass wall in a fish ladder; the camera is visible out of focus in o). Images d) to j) are from Mörrumsån, and images k) to l) are from Skeboån. Images m) and n) show corrupted frames that were retained in the dataset because corrupted images sometimes still contained detailed head regions of interest and increased variation in image quality. Many images in the full dataset did not contain a fish head, or did not contain a fish, but such negative/background images are not shown here.

The model was optimized using AdamW (Loshchilov and Hutter, 2019) with an initial learning rate of 0.003, a final learning-rate fraction of 0.1, weight decay of 0.01, and cosine learning-rate scheduling. A three-epoch warmup was used, with warmup momentum of 0.8 and a warmup bias learning rate of 0.1. Data augmentation included mosaic augmentation (Bochkovskiy et al., 2020), translation, scaling, and horizontal flipping, whereas mixup was disabled. Mosaic augmentation was disabled during the final 10 epochs to stabilize late-stage training.

The best checkpoint was used for validation and evaluation. Standard YOLO validation was performed on the validation split defined in the dataset configuration file, using an image size of 640 pixels and single-class evaluation. The extracted standard validation metrics were precision, recall, mAP50, and mAP50–95, following common object-detection evaluation practice (Lin et al., 2014).

Because the detector was used to generate head crops for downstream individual identification, an additional operational validation analysis was performed to evaluate how confidence-threshold choice affected crop generation. Validation images were resolved from the dataset configuration file, and the corresponding YOLO-format label files were read and converted from normalized centre-width-height coordinates to pixel-coordinate bounding boxes. Model predictions were first generated using a very low confidence threshold of 0.001 to retain nearly all candidate detections. During this prediction step, non-maximum suppression used an IoU threshold of 0.70 to remove overlapping duplicate boxes. Confidence thresholds were then applied retrospectively from 0.00 to 0.99 in increments of 0.01, with selected reporting thresholds of 0.50, 0.75, 0.90, and 0.95.

For each confidence threshold, predicted boxes were matched to annotated head boxes using greedy one-to-one matching. Predictions were sorted by confidence and matched to the unused ground-truth box with the highest IoU. A prediction was counted as a true positive when IoU was ≥ 0.50. Predictions that did not match any annotated head were counted as false boxes, and annotated heads without a matched prediction were counted as missed heads. For each threshold, the analysis recorded the number of validation images, annotated heads, predicted boxes, detected heads, missed heads, false boxes, precision, recall, F1 score, and mean IoU of matched detections.

The confidence threshold that produced the highest F1 score was identified from the threshold sweep. However, because the purpose of this model was to produce high-quality head crops for downstream identification, the final operating threshold was selected with emphasis on crop quality rather than maximum detection recall. Threshold-dependent performance was therefore reported to show the trade-off between retaining more annotated heads and producing fewer, higher-confidence crops.

#### 2.3.3 Individual re-identification model

Individual re-identification was performed using a deep metric-learning approach, in which cropped head images were mapped to normalized embedding vectors for identity comparison (Schroff et al., 2015). Images were organized by individual identity, with each identity represented by one folder. Only identity folders (n = 2,116) containing at least 20 images were considered trainable identities. Identity folders with fewer images (n = 895) were retained as non-training/unknown material for open-set evaluation rather than being used as training classes.

Before final model training, a short 3-epoch preliminary screening model was trained on the identity folders, comprising 194,178 images in total. Image embeddings were averaged within each identity folder, L2-normalized, and folder pairs were ranked by cosine similarity between these mean embeddings. This screening run used all trainable folders and an EfficientNetB0 model with an ArcFace head, using grayscale letterbox preprocessing without contrast-limited adaptive histogram equalization (CLAHE; Zuiderveld, 1994) or random augmentation. Its purpose was not to produce a final operational model, but to identify the pairs of individuals that were most difficult to separate, so that these hard pairs could be selected for final model training. By enriching the final training set with visually similar hard-negative identities, this selection step increased the likelihood that the small P–K batches contained informative identity contrasts, encouraging the model to learn fine-scale pattern differences among similar individuals despite limited GPU VRAM (Hermans et al., 2017). The final individual re-identification model was trained on 700 identity folders selected from a table created by the screening model. Folders were ranked by their involvement in high-similarity hard-pair rows, using the maximum cosine similarity to another trainable identity folder and the number of high-similarity rows in which the folder appeared. The 700 highest-ranked identity folders were used for supervised training. Most selected training identities originated from the Vindelälven material, with smaller contributions from Mörrumsån and Skeboån. All non-selected identity folders were kept outside model training and used as non-training identities for open-set and zero-shot evaluation.

All images were converted to grayscale before model input. For the final hard-pair model, CLAHE was applied, followed by letterbox resizing to 288 × 480 pixels. The resulting grayscale image was converted back to three channels so that it could be processed by an ImageNet-pretrained backbone. During training, augmentation was applied to improve robustness to natural variation in camera imagery. Augmentations included random gamma adjustment, brightness and contrast changes, additive Gaussian noise, Gaussian or motion blur, and small affine transformations including rotation, scaling, and translation. Horizontal flipping was not used.

The embedding model used an EfficientNetB0 backbone (Tan and Le, 2019) initialized with ImageNet weights (Deng et al., 2009). The convolutional backbone was followed by global average pooling, a 256-dimensional dense embedding layer, batch normalization (Ioffe and Szegedy, 2015), and L2 normalization. The network was trained using an ArcFace classification head with scale parameter 30 and angular margin 0.5 (Deng et al., 2019). Training used P–K sampling, with four identities and four images per identity in each batch, giving a batch size of 16 (Hermans et al., 2017). Optimization used AdamW (Loshchilov and Hutter, 2019) with a learning rate of 1 × 10⁻⁴ and weight decay of 1 × 10⁻⁴. Mixed-precision training was enabled to reduce memory use (Micikevicius et al., 2018). The final hard-pair model was trained for up to 25 epochs with model checkpointing; after training, the best checkpoint was reloaded before saving the embedding model and running evaluation.

For validation within the 700 trained identities, images were split into prototype/training and validation subsets using original video filename-based grouping rather than individual frames. The grouping key was defined as the part of the filename before the _f token, which marked the frame-number portion of extracted images. This was done to reduce leakage among near-duplicate frames from the same extracted video clip. Approximately 15% of available groups per identity were assigned to validation when more than one group was available. Because some identities were represented by only one filename group, not all trained identities contributed validation images. After splitting, the prototype/training set contained 78,273 images from 700 identities, and the grouped validation set contained 10,461 held-out images from 221 identities; 479 trained identities had no validation images.

After training, prototypes for the 700 trained identities were calculated by embedding the training images, averaging the resulting 256-dimensional embeddings within each identity, and L2-normalizing the mean vector, following the general prototype-based classification principle of representing each class by a mean embedding (Snell et al., 2017). Identification was then performed by comparing each query-image embedding with the relevant identity prototypes using cosine similarity (Schroff et al., 2015).

Threshold-based validation of known identities was performed using held-out validation images from the 700 trained identities. These validation images were not used to construct the trained-identity prototypes, but their identities were present in the training set. Images were split using a filename-grouped split, in which images sharing the same filename prefix before the frame-number token were kept together where possible to reduce leakage among near-duplicate frames. Approximately 15% of filename groups were assigned to validation, with the remaining groups used for prototype construction. Each validation image was compared with the trained-identity prototypes, and the nearest prototype was used as the candidate identity match. At the calibrated similarity threshold, performance was summarized as the proportion of known validation images accepted by the model and the accuracy of accepted known matches.

Open-set thresholding was evaluated using the same 700 trained-identity prototypes, but with images from non-training identities treated as unknown relative to the enrolled identities, following the open-set recognition setting in which test samples may come from classes not present during training (Scheirer et al., 2013). The non-training identities consisted of all identity folders not selected for training and were distinct from the held-out validation images from the trained identities. A similarity threshold was calibrated by comparing similarity scores from known validation images and a balanced sample of non-training images, with a target false-accept rate of approximately 1%. Final false-accept rates were then calculated on all non-training images and summarized both as an image-weighted false-accept rate and as folder-balanced false-accept rates across non-training folders.

Zero-shot retrieval was evaluated separately using the subset of non-training identity folders that contained at least 20 images. These 1,416 non-training identities were split internally into prototype and query images using the same filename-grouped splitting procedure as for the validation of the 700 hard-pair model. Approximately 15% of filename groups were assigned to the query subset and the remaining groups were used for prototype construction. For identity folders containing only one filename group, an image-level fallback split was used, assigning approximately 15% of images to the query subset and the remaining images to prototype construction while ensuring that both subsets contained at least one image. Prototypes were constructed from the prototype subset of each non-training identity, and query images were matched against prototypes from the same zero-shot identity set. Top-1, top-5, and top-10 retrieval accuracy were used to summarize zero-shot identification performance. This evaluation tested whether the learned embedding generalized to identities that were never used during model training.

### 2.4 Operational workflow

A practical software workflow is to stream camera footage from the NVR to a local analysis computer using RTSP and to screen the streams with the fish-presence detector. Rather than processing or storing all frames for downstream analysis, frames can be sampled at a reduced rate, for example one frame per second, to identify fish passages. When a fish is detected, a short video segment can be written with pre-and post-detection buffers to ensure that the complete passage event is retained. Camera identity and date-time information should be included in filenames, and camera clocks should be synchronized, because this spatiotemporal information is required for downstream capture–recapture analysis. Saved video clips can then be passed to the YOLOv8 ROI-extraction model to generate fish-head crops, and all ROI images from each clip can be placed in a single folder for downstream analysis. Detections near image borders can be excluded to avoid saving partial heads, and confidence-threshold filtering can be used to retain high-quality crops for the individual re-identification model. The extraction rate should be chosen according to study aims, storage capacity, and downstream processing needs, because full-frame-rate extraction can generate very large image datasets. The individual re-identification model is then used to merge folders containing the same individual and to split folders that contain more than one individual. After identities have been assigned and manually checked where necessary, camera identity and date-time information can be extracted from filenames to construct individual encounter histories. These encounter histories provide the individual-level data required for subsequent capture–recapture analysis.

## 3 Results

### 3.1 Machine vision pipeline

#### 3.1.1 Fish-presence detection model

The fish-presence detector was evaluated on a validation set containing 13,699 images, including 4,938 “fish” and 8,761 “no_fish” images. The best checkpoint achieved validation accuracy of 0.9918, loss of 0.0232, PR AUC of 0.9992, precision of 0.9873, recall of 0.9899, and ROC AUC of 0.9994. Threshold analysis showed that increasing the fish-probability threshold strongly reduced false positives while retaining high recall (Figure 3a,b). At a threshold of 0.98, the model produced 0 false positives among “no_fish” validation images and retained 94.3% “fish” recall, corresponding to 280 false negatives (Table 1).

**Figure 3.**
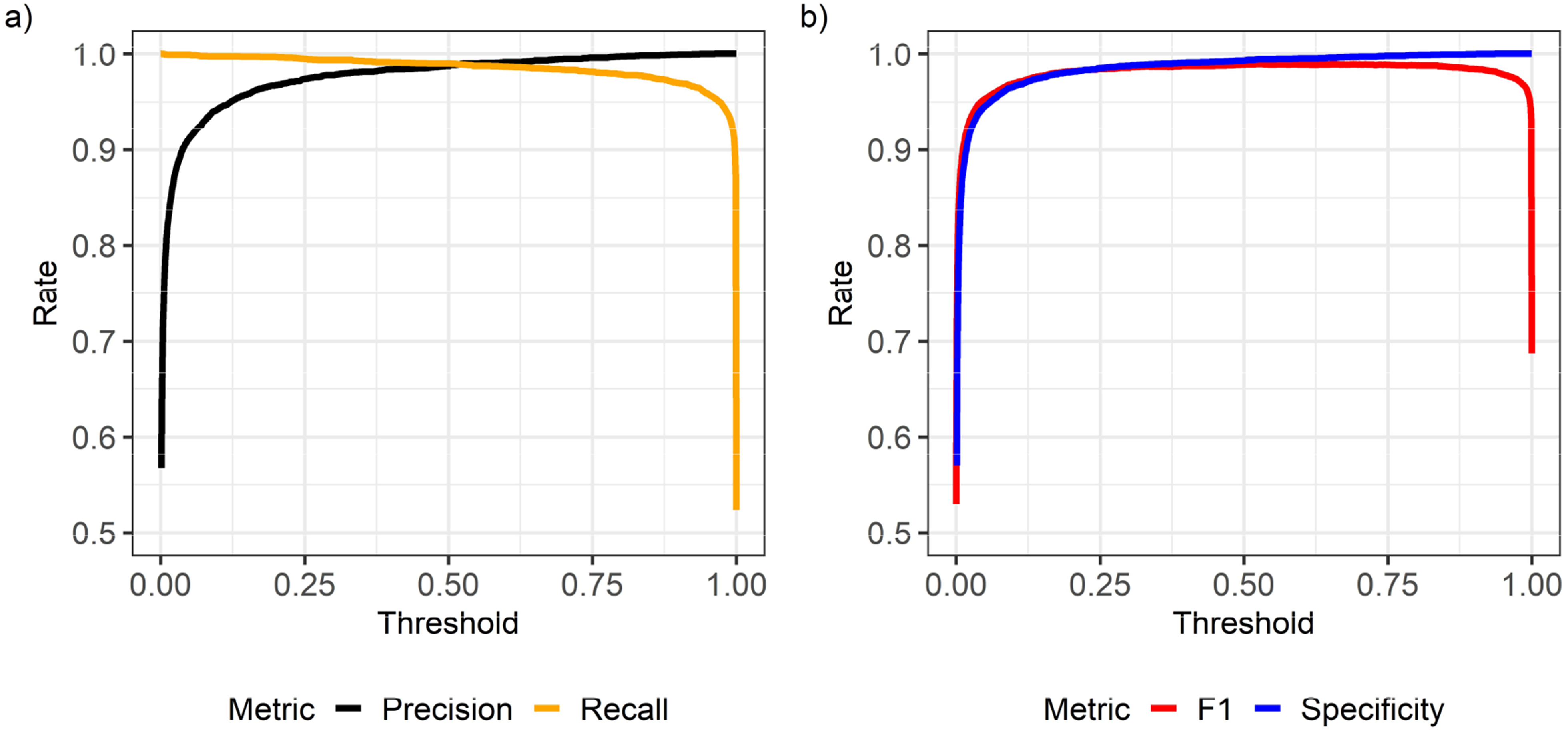
Threshold-dependent validation performance of the EfficientNetB0 fish-presence detector. Performance of the binary “fish” or “no_fish” classifier was evaluated across fish-probability thresholds from 0 to 1. Frames were classified as fish-positive when the predicted fish-probability was greater than or equal to the threshold. a) Precision and recall across thresholds, illustrating the trade-off between reducing false-positive fish detections and retaining true fish detections. b) Specificity and F1 score across thresholds, showing the model’s ability to reject “no_fish” frames and the overall balance between precision and recall. Metrics are shown for the validation set and were used to select conservative operating thresholds for screening video frames before region-of-interest cropping and individual-identification analysis.

**Table 1.** Threshold-specific validation performance of the EfficientNetB0 fish-presence detector. Performance of the binary “fish” or “no_fish” classifier was summarized at selected high fish-probability thresholds used to evaluate conservative video-frame screening. Frames were classified as fish-positive when the predicted fish-probability was greater than or equal to the threshold. Precision and recall are reported together with the number of false positives and false negatives in the validation set. False positives are “no_fish” images incorrectly classified as “fish”, whereas false negatives are “fish” images incorrectly classified as “no_fish”. These thresholds were evaluated to identify operating points that minimized false positives before downstream fish-head ROI detection and individual-identification analysis.

| Threshold | Precision | Recall | FP | FN |
| --- | --- | --- | --- | --- |
| 0.90 | 0.9990 | 0.9698 | 5 | 149 |
| 0.95 | 0.9996 | 0.9589 | 2 | 203 |
| 0.97 | 0.9998 | 0.9504 | 1 | 245 |
| 0.98 | 1.0000 | 0.9433 | 0 | 280 |
| 0.99 | 1.0000 | 0.9313 | 0 | 339 |

#### 3.1.2 Fish-head ROI detection model

The best YOLOv8n head-detection checkpoint was selected at epoch 68 based on validation mAP50–95. At this epoch, validation precision was 0.871, recall was 0.931, mAP50 was 0.967, and mAP50–95 was 0.867. Operational threshold analysis showed the expected trade-off between head-detection recall and crop selectivity (Figure 4a,b). At a confidence threshold of 0.90, the model produced 539 predicted boxes, of which 536 matched annotated heads and only 3 were false boxes, corresponding to a precision of 0.994, but with reduced recall because 896 annotated heads were missed (Table 2). This threshold was therefore selected as a conservative operating point for extracting high-quality head crops for downstream individual-identification analysis.

**Figure 4.**
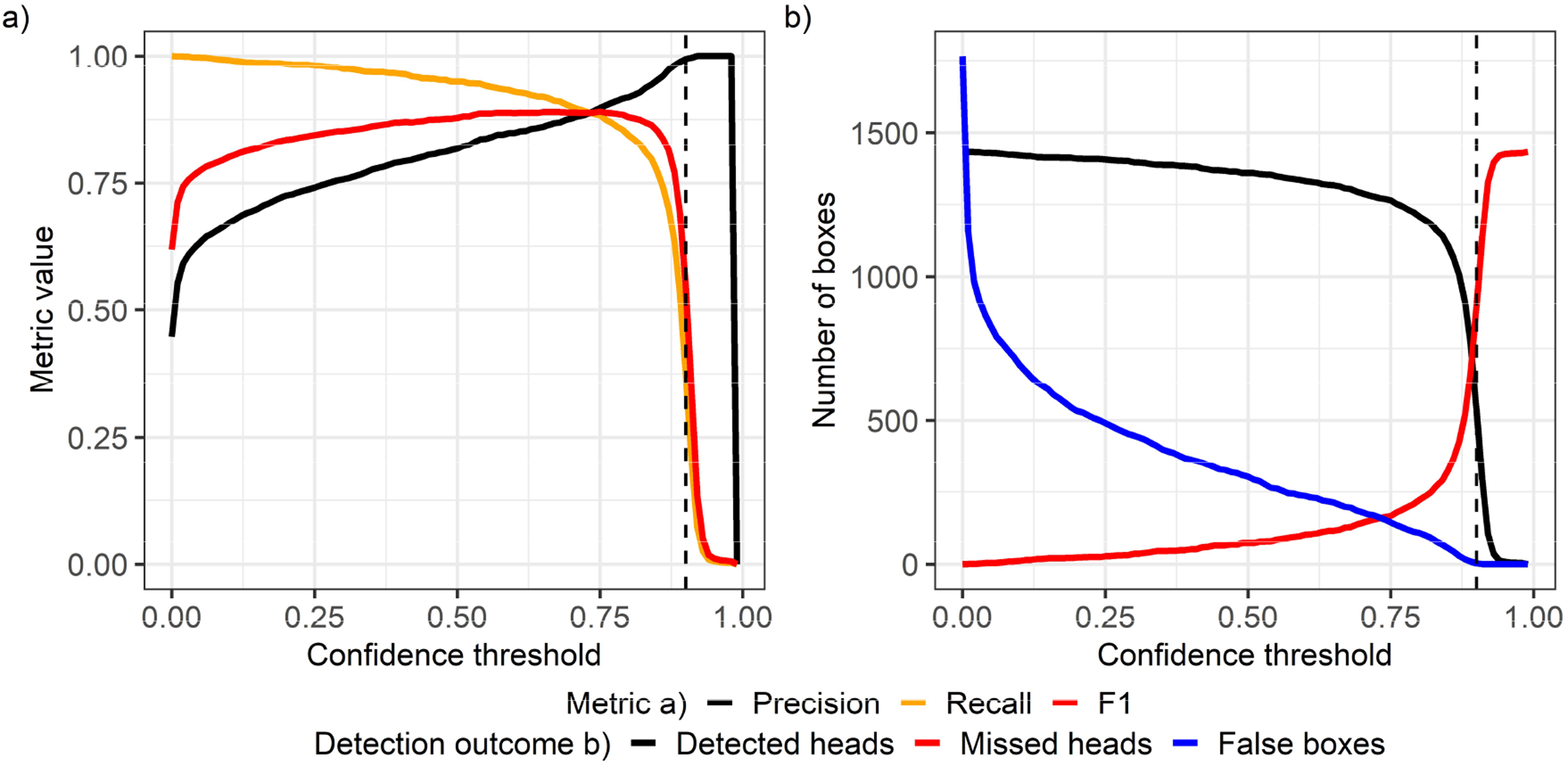
Threshold-dependent operational performance of the YOLOv8n fish-head detector on the validation set. Model performance was evaluated across confidence thresholds using one-to-one matching between predicted and annotated head boxes, with detections counted as correct when intersection-over-union was ≥ 0.50. Panel a) precision, recall, and F1 score across confidence thresholds. Panel b) number of detected heads, missed heads, and false predicted boxes across confidence thresholds. The dashed vertical line indicates the confidence threshold of 0.90 selected for conservative extraction of high-quality head crops for downstream individual-identification analysis.

**Table 2.**
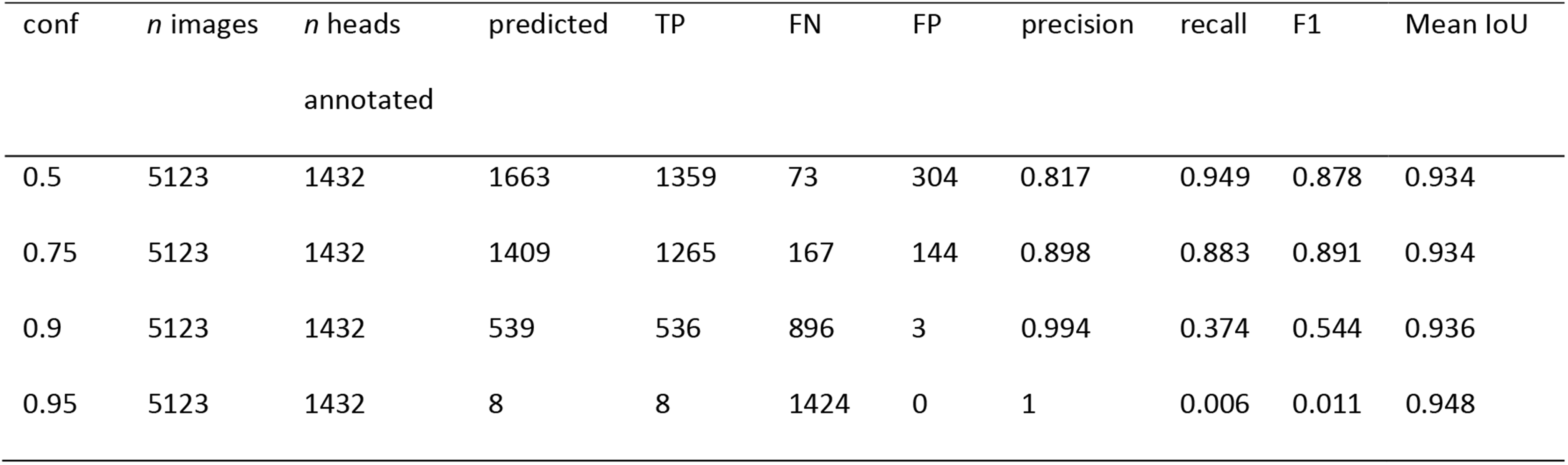
Operational validation performance of the YOLOv8n fish-head detector across selected confidence thresholds. Performance was evaluated on the validation set using a one-to-one matching rule between predicted and annotated head boxes, with detections considered correct when the intersection-over-union was ≥ 0.50. The table reports the number of validation images, annotated head instances, predicted boxes, detected heads, missed heads, false boxes, precision, recall, F1 score, and mean IoU of matched detections. Confidence thresholds were selected to illustrate the trade-off between head-detection recall and crop quality for downstream individual-identification analysis.

#### 3.1.3 Individual re-identification model

The final individual re-identification model was trained on 700 identities and evaluated using held-out images from trained identities, open-set images from non-training identities, and a separate zero-shot retrieval test. A similarity threshold of T = 0.514 was calibrated to target an approximately 1% false-accept rate (Figure 5). At this threshold, the model accepted 91.05% of held-out images from trained identities, and 99.80% of accepted known matches were assigned to the correct identity. When all remaining non-training identities were treated as unknown, the image-weighted false-accept rate was 0.81%, while the folder-balanced false-accept rate was 1.17% across all non-training folders and 0.85% among folders containing at least 20 images. In the zero-shot evaluation, which tested identities not used during model training, the model achieved 99.17% top-1 accuracy, 99.68% top-5 accuracy, and 99.80% top-10 accuracy across 1,416 non-training identities and 22,175 query images. These results indicate that the embedding model separated known and unknown identities well and generalized strongly to identities excluded from training (Table 3). Cosine similarities between selected individual pairs are illustrated in Figure 6 to show examples of a range of visually similar and visually distinct identities in the learned embedding space.

**Figure 5.**
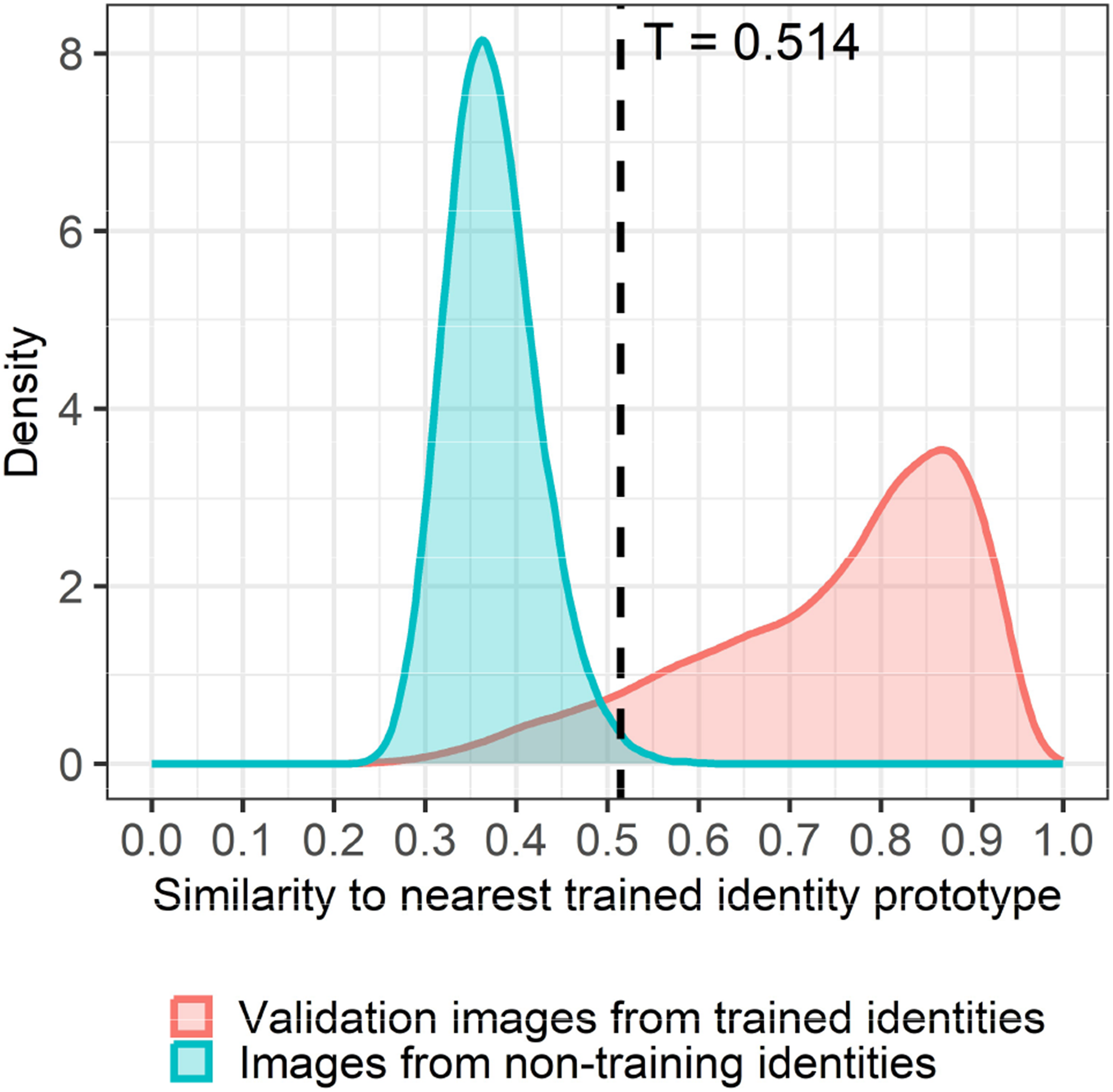
Distribution of image-level similarity scores for trained and non-training identities. Density distributions show the similarity of each image to the nearest trained identity prototype. Validation images from trained identities were held-out images from the 700 identities used to train the model (n = 10,461 images), whereas images from non-training identities were from identity folders excluded from model training (n = 112,904 images). The dashed vertical line shows the calibrated similarity threshold, T = 0.514, used to accept or reject a candidate match. Images from trained identities falling below the threshold are rejected, whereas images from non-training identities exceeding the threshold represent false accepts. The threshold was calibrated to target an approximately 1% false-accept rate.

**Figure 6.**
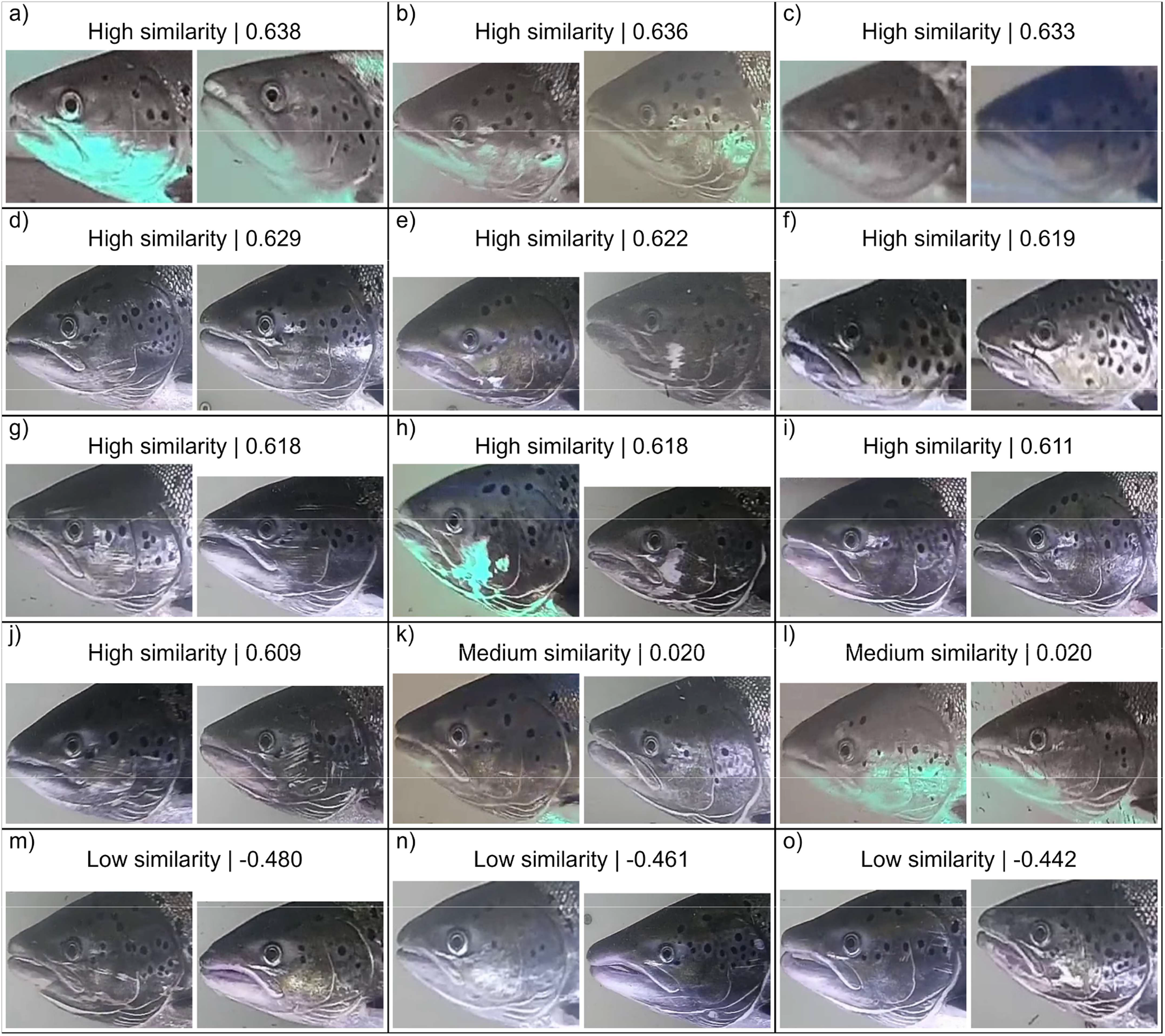
Fifteen pairs of identity prototypes are shown to illustrate the range of visual similarity in the embedding space, where each image shows a unique individual fish. The first ten panels a) to j) show the highest similarity prototype pairs, representing identities that the model judged to be visually close, whereas panels k) and l) show pairs of medium similarity and the final three panels m) to o) show low similarity pairs for contrast. Each panel contains one image per identity folder, selected as the image with the highest YOLO detection confidence; hence the displayed images are illustrative, whereas the reported similarity is calculated between folder-level identity prototypes. The value above each pair is the cosine similarity between the two identity prototypes, where higher values indicate greater similarity in the learned embedding space.

**Table 3.** Performance of the individual re-identification model. Summary of trained-identity validation, open-set false-accept performance, and zero-shot retrieval performance for the final individual re-identification model. The model was trained on 700 identities. Open-set performance was evaluated by treating all remaining non-training identities as unknown relative to the trained identity set. The similarity threshold, T, was calibrated to target an approximately 1% false-accept rate. Image-weighted false-accept rate was calculated across all images from non-training identities, whereas folder-balanced false-accept rate was calculated by averaging false-accept rates across non-training ID folders. Zero-shot retrieval was evaluated separately using non-training identities with at least 20 images. For this zero-shot evaluation, 83,265 prototype images were used to construct identity prototypes, and 22,175 held-out query images were matched against those prototypes to calculate top-1, top-5, and top-10 retrieval accuracy.

| Metric | Value |
| --- | --- |
| Trained identities | 700 |
| Non-training images tested as unknown, all ID folders | 112,904 |
| Non-training ID folders tested as unknown | 2,311 |
| Threshold T | 0.514 |
| Image-weighted false-accept rate | 0.81% |
| Folder-balanced false-accept rate, all non-training ID folders | 1.17% |
| Folder-balanced false-accept rate, non-training ID folders $\geq 20$ images | 0.85% |
| Known accept rate | 91.05% |
| Accuracy among accepted known matches | 99.80% |
| Zero-shot identities, non-training ID folders $\geq 20$ images | 1,416 |
| Zero-shot query images | 22,175 |
| Zero-shot prototype images | 83,265 |
| Zero-shot top-1 accuracy | 99.17% |
| Zero-shot top-5 accuracy | 99.68% |
| Zero-shot top-10 accuracy | 99.80% |

## 4 Discussion

The machine-vision pipeline presented here outlines a cohesive workflow for monitoring salmon and sea trout spawning runs in riverine habitats within a two-camera-station capture–recapture design. A major strength of probability-based monitoring, such as capture–recapture modelling, is that the unobserved part of the population can be estimated from observed capture and recapture data by explicitly accounting for detection probability (Otis et al., 1978; Kéry and Royle, 2021). Consequently, only a subset of the population needs to be detected, and the monitoring setup does not require major interventions such as dams, blockades, or fish ladders that force all upstream-migrating fish through a single passage. This makes the approach relevant to two-station salmonid monitoring designs, where downstream detections and upstream re-identifications can be used to estimate detection probability and abundance (Schwarz and Dempson, 1994; Mäntyniemi and Romakkaniemi, 2002). The pipeline may therefore also be useful in unexploited and free-flowing river systems, provided that enough individuals are detected and recaptured. In such systems, cameras could be deployed at locations where fish naturally aggregate, such as river narrows and deep pools, or combined with minor guiding structures that partially direct fish toward the camera field of view. The proposed method therefore has potential for estimating salmon and trout population size in both exploited and less-modified river systems.

Monitoring complete spawning and upstream-migration periods remains challenging because salmonid run timing can extend over several months and vary among stocks, rivers, and years (Klemetsen et al., 2003; Weedop et al., 2026). Long-duration monitoring therefore requires stable camera operation, sufficient storage, reliable power and network connections, and periodic maintenance. In the workflow presented here, NVR-based recording combined with automated fish detection and clip extraction provides a practical way to reduce manual video review while retaining the individual-level information needed for population estimation. The main practical challenge is therefore obtaining sufficient high-quality detections and recaptures; as computational capacity continues to increase, field data collection is likely to become a greater constraint than data analysis.

This embedding-based approach could be extended to other monitoring settings and species with stable, individually distinctive markings, including fish photographed during capture-based or angler-based sampling (Karlsson and Kari, 2020). In such settings, each image can often be linked to a single handled individual, reducing the multi-individual ambiguity present in continuous video and allowing more standardized image acquisition. Future improvements could also come from larger training batches and more extensive training data, as GPU memory limits the number of identities and images included in each P–K batch (Hermans et al., 2017). In addition, promptable segmentation models such as SAM or SAM 2 could be evaluated as intermediate steps between head detection and individual identification, but their computational cost would need to be weighed against any improvement in crop quality and identification accuracy (Kirillov et al., 2023; Ravi et al., 2024).

Overall, the results show that machine-vision-based individual identification can generate the encounter histories required for capture–recapture monitoring of salmonid spawning runs. As computing power and vision models continue to improve, a major future challenge is likely to be field data collection: obtaining sufficient high-quality images, detections, and recaptures under realistic environmental conditions.

## Acknowledgements

I thank Vattenfall R&D for providing video data from the fish passage facility at Stornorrfors in the Vindelälven, and Sveaskog Kronolaxfiske Mörrumsån for maintaining the camera and monitoring equipment in Mörrumsån. I am also grateful for support during the Energiforsk project application and early project development.

## Data availability

Processed datasets, evaluation outputs, and scripts required to reproduce the reported figures and tables, together with the complete image datasets and additional scripts used for model training and evaluation, will be added to a public Zenodo record before publication of the peer-reviewed article.

## Declaration of generative AI

During the preparation of this work, the author used ChatGPT by OpenAI for language editing, manuscript-structure suggestions, drafting support, and assistance with writing and debugging analysis scripts. Generative AI was not used to annotate images, assign fish identities, create training labels, or generate or alter primary image data used in the machine-vision pipeline. After using this tool, the author reviewed, edited, and verified the outputs as needed and takes full responsibility for the content of this manuscript.

## References

1. Abadi, M., Barham, P., Chen, J., Chen, Z., Davis, A., Dean, J., … Zheng, X. (2016). TensorFlow: A system for large-scale machine learning. In 12th USENIX Symposium on Operating Systems Design and Implementation (OSDI 16) (pp. 265–283). USENIX Association. Retrieved from https://www.usenix.org/conference/osdi16/technical-sessions/presentation/abadi

2. Atlas, W. I., Ma, S., Chou, Y. C., Connors, K., Scurfield, D., Nam, B., … Liu, J. (2023). Wild salmon enumeration and monitoring using deep learning empowered detection and tracking. Frontiers in Marine Science, 10, 1200408. 10.3389/fmars.2023.1200408

3. Bishop, C. M. (2006). Pattern recognition and machine learning. New York, NY: Springer.

4. Bochkovskiy, A., Wang, C.-Y., & Liao, H.-Y. M. (2020). *YOLOv4: Optimal speed and accuracy of object detection* [Preprint]. arXiv. 10.48550/arXiv.2004.10934

5. Bowman, J. C., Fedus, A. L., Fox, M. G., & Raby, G. D. (2026). How useful is underwater video as a fisheries assessment tool in temperate freshwater ecosystems? Transactions of the American Fisheries Society, 155(2), 200–213. 10.1093/tafafs/vnaf057

6. Charbonneau, J. A., Connors, K., Atkinson, C., Hertz, E., Connors, B., & Moore, J. W. (2025). Hyperstability in an inland recreational fishery: Are catch-per-unit-effort data masking the magnitude of steelhead declines? Transactions of the American Fisheries Society, 154(4), 339–351. 10.1093/tafafs/vnaf023

7. Chollet, F. (2015). *Keras* [Computer software]. Retrieved from https://github.com/keras-team/keras

8. Chollet, F. (2017). Deep learning with Python. Shelter Island, NY: Manning Publications.

9. Christin, S., Hervet, É., & Lecomte, N. (2019). Applications for deep learning in ecology. Methods in Ecology and Evolution, 10(10), 1632–1644. 10.1111/2041-210X.13256

10. Deng, J., Dong, W., Socher, R., Li, L.-J., Li, K., & Fei-Fei, L. (2009). ImageNet: A large-scale hierarchical image database. In 2009 IEEE Conference on Computer Vision and Pattern Recognition. IEEE. 10.1109/CVPR.2009.5206848

11. Deng, J., Guo, J., Xue, N., & Zafeiriou, S. (2019). ArcFace: Additive angular margin loss for deep face recognition. In Proceedings of the IEEE/CVF Conference on Computer Vision and Pattern Recognition, 4690–4699. https://openaccess.thecvf.com/content_CVPR_2019/papers/Deng_ArcFace_Additive_Angular_Margin_Loss_for_Deep_Face_Recognition_CVPR_2019_paper.pdf

12. Erisman, B. E., Allen, L. G., Claisse, J. T., Pondella, D. J., Miller, E. F., & Murray, J. H. (2011). The illusion of plenty: Hyperstability masks collapses in two recreational fisheries that target fish spawning aggregations. Canadian Journal of Fisheries and Aquatic Sciences, 68(10), 1705–1716. 10.1139/f2011-090

13. Fawcett, T. (2006). An introduction to ROC analysis. Pattern Recognition Letters, 27(8), 861–874. 10.1016/j.patrec.2005.10.010

14. HELCOM. (2011). Salmon and sea trout populations and rivers in the Baltic Sea—HELCOM assessment of salmon (Salmo salar) and sea trout (Salmo trutta) populations and habitats in rivers flowing to the Baltic Sea (Baltic Sea Environment Proceedings No. 126). Retrieved from https://helcom.fi/wp-content/uploads/2019/08/BSEP126A.pdf

15. Hermans, A., Beyer, L., & Leibe, B. (2017). *In defense of the triplet loss for person re-identification* [Preprint]. arXiv. 10.48550/arXiv.1703.07737

16. Ioffe, S., & Szegedy, C. (2015). Batch normalization: Accelerating deep network training by reducing internal covariate shift. In Proceedings of the 32nd International Conference on Machine Learning 37, 448–456. Proceedings of Machine Learning Research. http://proceedings.mlr.press/v37/ioffe15.pdf

17. Jocher, G., Chaurasia, A., & Qiu, J. (2023). *Ultralytics YOLOv8* (Version 8.0.0) [Computer software]. Retrieved from https://github.com/ultralytics/ultralytics

18. Karanth, K. U. (1995). Estimating tiger *Panthera tigris* populations from camera-trap data using capture–recapture models. Biological Conservation, 71(3), 333–338. 10.1016/0006-3207(94)00057-W

19. Karlsson, K. (2024). A hands-on guide to use network video recorders, internet protocol cameras, and deep learning models for dynamic monitoring of trout and salmon in small streams. Ecology and Evolution, 14(5), e11246. 10.1002/ece3.11246

20. Karlsson, K., Andersson, H. C., & Sundblad, G. (2025). Estimating northern pike population dynamics and capture probability by recreational angling using spatial capture–recapture models in a Baltic Sea spawning area. Transactions of the American Fisheries Society, 155(2), 160–176. 10.1093/tafafs/vnaf053

21. Karlsson, K., & Kari, E. (2020). Recreational anglers as citizen scientists can provide data to estimate population size of pike, *Esox lucius*. Fisheries Management and Ecology, 27(4), 367–380. 10.1111/fme.12419

22. Kazyak, D. C., Flowers, A. M., Hostetter, N. J., Madsen, J. A., Breece, M. W., Higgs, A., … Fox, D. A. (2020). Integrating side-scan sonar and acoustic telemetry to estimate the annual spawning run size of Atlantic sturgeon in the Hudson River. Canadian Journal of Fisheries and Aquatic Sciences, 77(6), 1038–1048. 10.1139/cjfas-2019-0398

23. Kéry, M., & Royle, J. A. (2016). Applied hierarchical modeling in ecology: Analysis of distribution, abundance and species richness in R and BUGS: Vol. 1. Prelude and static models. London, England: Academic Press.

24. Kéry, M., & Royle, J. A. (2021). Applied hierarchical modeling in ecology: Analysis of distribution, abundance and species richness in R and BUGS: Vol. 2. Dynamic and advanced models. London, England: Academic Press.

25. Kingma, D. P., & Ba, J. (2015). Adam: A method for stochastic optimization. In International Conference on Learning Representations. 10.48550/arXiv.1412.6980

26. Kirillov, A., Mintun, E., Ravi, N., Mao, H., Rolland, C., Gustafson, L., … Girshick, R. (2023). Segment Anything. In Proceedings of the IEEE/CVF International Conference on Computer Vision (pp. 4015–4026). 10.1109/ICCV51070.2023.00371

27. Klemetsen, A., Amundsen, P.-A., Dempson, J. B., Jonsson, B., Jonsson, N., O’Connell, M. F., & Mortensen, E. (2003). Atlantic salmon *Salmo salar* L., brown trout *Salmo trutta* L. and Arctic charr *Salvelinus alpinus* (L.): A review of aspects of their life histories. Ecology of Freshwater Fish, 12(1), 1–59. 10.1034/j.1600-0633.2003.00010.x

28. Krizhevsky, A., Sutskever, I., & Hinton, G. E. (2012). ImageNet classification with deep convolutional neural networks. Advances in Neural Information Processing Systems, 25, 1097–1105. https://proceedings.neurips.cc/paper_files/paper/2012/file/c399862d3b9d6b76c8436e924a68c45b-Paper.pdf

29. Lin, T.-Y., Maire, M., Belongie, S., Bourdev, L., Girshick, R., Hays, J., … Zitnick, C. L. (2014). Microsoft COCO: Common objects in context. In D. Fleet, T. Pajdla, B. Schiele, & T. Tuytelaars (Eds.), Computer vision—ECCV 2014 (Lecture Notes in Computer Science, Vol. 8693, pp. 740–755). Cham, Switzerland: Springer. 10.1007/978-3-319-10602-1_48

30. Loshchilov, I., & Hutter, F. (2019). Decoupled weight decay regularization. In International Conference on Learning Representations. 10.48550/arXiv.1711.05101

31. Micikevicius, P., Narang, S., Alben, J., Diamos, G., Elsen, E., Garcia, D., … Wu, H. (2018). Mixed precision training. In International Conference on Learning Representations. 10.48550/arXiv.1710.03740

32. Mäntyniemi, S., & Romakkaniemi, A. (2002). Bayesian mark–recapture estimation with an application to a salmonid smolt population. Canadian Journal of Fisheries and Aquatic Sciences, 59(11), 1748– 1758. 10.1139/f02-146

33. Norouzzadeh, M. S., Nguyen, A., Kosmala, M., Swanson, A., Palmer, M. S., Packer, C., & Clune, J. (2018). Automatically identifying, counting, and describing wild animals in camera-trap images with deep learning. Proceedings of the National Academy of Sciences of the United States of America, 115(25), E5716–E5725. 10.1073/pnas.1719367115

34. Otis, D. L., Burnham, K. P., White, G. C., & Anderson, D. R. (1978). Statistical inference from capture data on closed animal populations. Wildlife Monographs, 62, 3–135. https://www.jstor.org/stable/3830650

35. Raabe, J. K., Gardner, B., & Hightower, J. E. (2014). A spatial capture–recapture model to estimate fish survival and location from linear continuous monitoring arrays. Canadian Journal of Fisheries and Aquatic Sciences, 71(1), 120–130. 10.1139/cjfas-2013-0198

36. Ravi, N., Gabeur, V., Hu, Y.-T., Hu, R., Ryali, C., Ma, T., … Feichtenhofer, C. (2024). *SAM 2: Segment Anything in images and videos* [Preprint]. arXiv. 10.48550/arXiv.2408.00714

37. Redmon, J., Divvala, S., Girshick, R., & Farhadi, A. (2016). You only look once: Unified, real-time object detection. In Proceedings of the IEEE Conference on Computer Vision and Pattern Recognition (pp. 779–788).

38. Roser, P., Radinger, J., Feldhege, F., Braun, M., & Arlinghaus, R. (2025). Getting scarce and lure shy: Impacts of recreational fishing on coastal northern pike (*Esox lucius*) abundance, size structure and vulnerability to angling. Fisheries Management and Ecology, 32(3), e12769. 10.1111/fme.12769

39. Royle, J. A., Fuller, A. K., & Sutherland, C. (2018). Unifying population and landscape ecology with spatial capture–recapture. Ecography, 41(3), 444–456. 10.1111/ecog.03170

40. Saito, T., & Rehmsmeier, M. (2015). The precision–recall plot is more informative than the ROC plot when evaluating binary classifiers on imbalanced datasets. PLOS ONE, 10(3), e0118432. 10.1371/journal.pone.0118432

41. Scheirer, W. J., de Rezende Rocha, A., Sapkota, A., & Boult, T. E. (2013). Toward open set recognition. IEEE Transactions on Pattern Analysis and Machine Intelligence, 35(7), 1757–1772. 10.1109/TPAMI.2012.256

42. Schneider, S., Taylor, G. W., Linquist, S., & Kremer, S. C. (2019). Past, present and future approaches using computer vision for animal re-identification from camera trap data. Methods in Ecology and Evolution, 10(4), 461–470. 10.1111/2041-210X.13133

43. Schroff, F., Kalenichenko, D., & Philbin, J. (2015). FaceNet: A unified embedding for face recognition and clustering. In Proceedings of the IEEE Conference on Computer Vision and Pattern Recognition, 815–823. https://www.cv-foundation.org/openaccess/content_cvpr_2015/papers/Schroff_FaceNet_A_Unified_2015_CVPR_paper.pdf

44. Schwarz, C. J., & Dempson, J. B. (1994). Mark-recapture estimation of a salmon smolt population. Biometrics, 50(1), 98–108. 10.2307/2533200

45. Snell, J., Swersky, K., & Zemel, R. S. (2017). Prototypical networks for few-shot learning. Advances in Neural Information Processing Systems, 30. https://proceedings.neurips.cc/paper_files/paper/2017/file/cb8da6767461f2812ae4290eac7cbc42-Paper.pdf

46. Soom, J., Pattanaik, V., Leier, M., & Tuhtan, J. A. (2022). Environmentally adaptive fish or no-fish classification for river video fish counters using high-performance desktop and embedded hardware. Ecological Informatics, 72, 101817. 10.1016/j.ecoinf.2022.101817

47. Srivastava, N., Hinton, G., Krizhevsky, A., Sutskever, I., & Salakhutdinov, R. (2014). Dropout: A simple way to prevent neural networks from overfitting. Journal of Machine Learning Research, 15(56), 1929–1958. http://jmlr.org/papers/v15/srivastava14a.html

48. Steenweg, R., Hebblewhite, M., Kays, R., Ahumada, J. A., Fisher, J. T., Burton, C., … Rich, L. N. (2017). Scaling-up camera traps: Monitoring the planet’s biodiversity with networks of remote sensors. Frontiers in Ecology and the Environment, 15(1), 26–34. 10.1002/fee.1448

49. Tan, M., & Le, Q. V. (2019). EfficientNet: Rethinking model scaling for convolutional neural networks. In Proceedings of the 36th International Conference on Machine Learning, 97, 6105–6114. Proceedings of Machine Learning Research http://proceedings.mlr.press/v97/tan19a/tan19a.pdf

50. Vidal, M., Wolf, N., Rosenberg, B., Harris, B. P., & Mathis, A. (2021). Perspectives on individual animal identification from biology and computer vision. Integrative and Comparative Biology, 61(3), 900– 916. 10.1093/icb/icab107

51. Weedop, D., Womer, J. D., Ziller, J. S., & Murphy, C. A. (2026). Timing is everything: Drivers of upstream movement of fishes. Transactions of the American Fisheries Society, 155(4), 368–383. 10.1093/tafafs/vnag008

52. Weinstein, B. G. (2018). A computer vision for animal ecology. Journal of Animal Ecology, 87(3), 533–545. 10.1111/1365-2656.12780

53. Würsig, B., & Jefferson, T. A. (1990). Methods of photoidentification for small cetaceans. Reports of the International Whaling Commission, 12(Special Issue), 43–52.

54. Yesharim, M., Perl, R. B., Roll, U., Gafny, S., Geffen, E., & Ram, Y. (2026). Near-perfect photo-ID of the Hula painted frog with zero-shot deep local-feature matching. Ecological Informatics, 98, 103942. 10.1016/j.ecoinf.2026.103942

55. Zuiderveld, K. (1994). Contrast limited adaptive histogram equalization. In P. S. Heckbert (Ed.), Graphics gems IV (pp. 474–485). San Diego, CA: Academic Press.

